# Genomic and chromatin features shaping meiotic double-strand break formation and repair in mice

**DOI:** 10.1101/131748

**Authors:** Shintaro Yamada, Seoyoung Kim, Sam E. Tischfield, Julian Lange, Maria Jasin, Scott Keeney

**Affiliations:** Molecular Biology Program, Memorial Sloan Kettering Cancer Center, New York, NY, USA; Howard Hughes Medical Institute, Memorial Sloan Kettering Cancer Center, New York, NY, USA; Tri-Institutional Training Program in Computational Biology and Medicine, Weill Cornell Medical College, New York, NY, USA; Developmental Biology Program, Memorial Sloan Kettering Cancer Center, New York, NY, USA

**Keywords:** SPO11, meiosis, recombination, DNA double-strand breaks, resection, PRDM9, chromatin, repetitive elements

## Abstract

The SPO11-generated DNA double-strand breaks (DSBs) that initiate meiotic recombination occur non-randomly across genomes, but mechanisms shaping their distribution and repair remain incompletely understood. Here, we expand on recent studies of nucleotide-resolution DSB maps in mouse spermatocytes. We find that trimethylation of histone H3 lysine 36 around DSB hotspots is highly correlated, both spatially and quantitatively, with trimethylation of H3 lysine 4, consistent with coordinated formation and action of both PRDM9-dependent histone modifications. In contrast, the DSB-responsive kinase ATM contributes independently of PRDM9 to controlling hotspot activity, and combined action of ATM and PRDM9 can explain nearly two-thirds of the variation in DSB frequency between hotspots. DSBs were modestly underrepresented in most repetitive sequences such as segmental duplications and transposons. Nonetheless, numerous DSBs form within repetitive sequences in each meiosis and some classes of repeats are preferentially targeted. Implications of these findings are discussed for evolution of PRDM9 and its role in hybrid strain sterility in mice. Finally, we document the relationship between mouse strain-specific DNA sequence variants within PRDM9 recognition motifs and attendant differences in recombination outcomes. Our results provide further insights into the complex web of factors that influence meiotic recombination patterns.

## Introduction

Cells undergoing meiosis inflict DNA double-strand breaks (DSBs) at many places across the genome to initiate recombination, which physically links homologous chromosomes to promote their accurate segregation. These DSBs occur preferentially (but not exclusively) within highly localized regions called hotspots.^1^ The non-random distribution of DSBs governs the evolution and diversity of eukaryotic genomes. Furthermore, failure to properly form and repair meiotic DSBs results in gametes with chromosome structure alterations or aneuploidy, which can lead to developmental disorders.^2,3^ An important challenge has been to understand the complex interplay of multiple factors that shape this DSB landscape over size scales ranging from single nucleotides to whole chromosomes.^4-6^

Meiotic DSBs are formed by dimers of the conserved topoisomerase-like protein SPO11 via a transesterase reaction that links a SPO11 molecule to each 5′ end of the broken DNA.^7,8^ DNA nicks then release SPO11 covalently bound to short oligonucleotides (SPO11 oligos).^9^ 5′→3′ resection generates a single-stranded DNA (ssDNA) tail at each DSB end.^10-13^ This ssDNA becomes coated with strand-exchange proteins DMC1 and RAD51 and searches for homologous DNA as a repair template.^14-17^ DSB repair is completed as a crossover (reciprocal exchange of chromosome arms that flank the repair site) or a noncrossover.^18^

Among many levels of mammalian DSB landscape organization, hotspot control by the histone methyltransferase PRDM9 has been the most extensively studied. In mice and humans, PRDM9 is a major determinant of hotspot locations via its sequence-specific, multi-zinc-finger DNA-binding domain, the specificity of which evolves rapidly and is highly polymorphic in populations.^19,20^ Genome-wide hotspot distributions in these two organisms have been examined by mapping DMC1-bound ssDNA or PRDM9-dependent histone H3 lysine 4 trimethylation (H3K4me3).^21-25^ However, constraints on spatial resolution of these maps left questions unanswered about fine-scale DSB patterns, especially at the sub-hotspot level. We recently overcame this issue by sequencing SPO11 oligos purified from mouse testes.^26^ SPO11-oligo mapping provided quantitative DSB landscapes at nucleotide resolution, with low background and high dynamic range. The fine-scale maps revealed previously invisible spatial features of DSBs and relationships between SPO11, PRDM9 and methylated nucleosomes. Here, we combine these SPO11-oligo data with other published genome-wide data to further explore genomic features that influence DSB formation and repair in mice.

## Results and Discussion

### Relationships of DSB patterns with the trimethylation of H3K36 and H3K4

Studies of DSB and H3K4me3 distributions have shown that DSBs are targeted by PRDM9 to genomic regions that are recognized by the PRDM9 zinc finger DNA-binding domain and subject to local histone methylation by its PR/SET methyltransferase domain.^21,22,24,27-29^ Previous work showed that the mean H3K4me3 signal oscillates around DSB hotspots, with immediately adjacent nucleosomes showing stronger H3K4me3 signal than nucleosomes further away (**Fig. 1A**).^22,26^ However, PRDM9 also trimethylates histone H3 on lysine 36 *in vitro* and *in vivo*, ^24,30-32^ so we examined this modification as well using published H3K36me3 chromatin immunoprecipitation (ChIP) data.^32^

**Figure 1.**
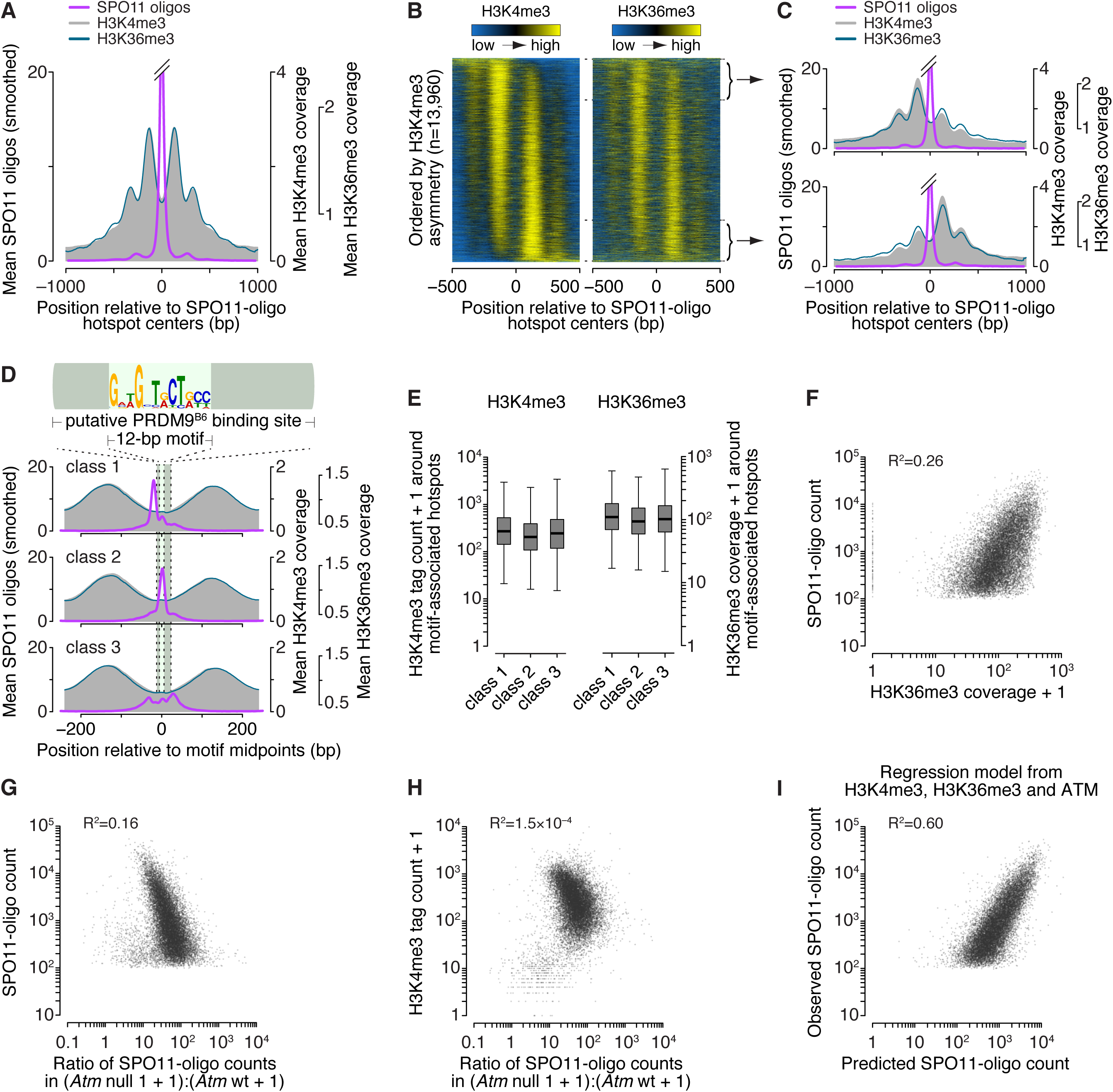
Spatial relationships between H3K36 trimethylation and DSBs. (A) H3K36me3 (data from ref.^32^) has a similar profile as H3K4me3 (data from ref.^22^) around SPO11-oligo hotspots. Data were locally normalized by dividing the signal at each base pair (bp) by the mean signal within each 2,001-bp window, then were averaged across hotspots. The SPO11-oligo profile was smoothed with a 51-bp Hann filter. (B) H3K36me3 signal is often highly asymmetric around hotspots in a manner similar to H3K4me3. Heat maps (data in 5-bp bins after local normalization) were ordered according to H3K4me3 asymmetry. Because data in each hotspot were locally normalized, color-coding reflects the local spatial pattern, not relative signal strength between hotspots. (C) Similar asymmetric patterns between H3K4me3 and H3K36me3 at SPO11-oligo hotspots. Each panel shows the mean of locally normalized profiles (51-bp Hann filter for SPO11-oligo data) across the 20% of hotspots with the most asymmetric H3K4me3 patterns (left > right in top panel; right > left in bottom panel). (D) In three classes of PRDM9 motifs previously defined according to local SPO11-oligo pattern,^26^ H3K36me3 patterns are similar. The sequence logo shows the 12-bp core PRDM9 motif identified by MEME within SPO11-oligo hotspots;^26^ the light gray bar denotes the larger 36-bp segment of DNA thought to be bound by PRDM9.^21,29^ SPO11-oligo profiles were smoothed with a 15-bp Hann filter. (E) Similar H3K36me3 signal strength for hotspots in each of the three PRDM9 motif classes, as observed for H3K4me3. H3K4me3 tag counts and H3K36me3 coverage were summed in 1,001-bp windows around hotspot centers. In the box plots, thick horizontal lines indicate medians, box edges show the 25th and 75th percentiles, and whiskers indicate lowest and highest values within 1.5-fold of the interquartile range; outliers are not shown. A value of 1 was added to each hotspot to permit plotting of hotspots with no H3K4me3 or H3K36me3 signal. (F) H3K36me3 is an imperfect predictor of DSB frequency. SPO11-oligo counts and H3K36me3 coverage were summed in 1,001-bp windows around hotspot centers. One H3K36me3 count was added to each hotspot to permit plotting of hotspots with no H3K36me3 signal. (G–H) The effect of ATM deficiency on hotspot activity is independent of H3K4me3 levels. SPO11-oligo counts in B6, *Atm* null and *Atm* wt and H3K4me3 tag counts were summed in 1,001-bp windows around B6 hotspot centers. The ratio of SPO11-oligo counts in *Atm* null to *Atm* wt was plotted against SPO11-oligo counts in B6 (G) or H3K4me3 counts (H). One count was added to each hotspot in *Atm* null and *Atm* wt SPO11-oligo data and H3K4me3 data to permit plotting of hotspots with no signal. (I) Fit of a multiple regression model predicting SPO11-oligo counts in hotspots from H3K4me3, H3K36me3, and *Atm* null:*Atm* wt ratio (**Table S1**). Panels depicting SPO11-oligo and H3K4me3 ChIP data were adapted from ref ^26^ with permission. In panels showing H3K36me3 signal around SPO11-oligo hotspots and PRDM9 motifs, data were plotted starting from values of 0.56 to facilitate side-by-side comparison with H3K4me3 data. All R^2^ values were determined by linear regression on log-transformed data.

When averaged around SPO11-oligo hotspot centers, the H3K36me3 pattern was strikingly similar to that of H3K4me3 (**Fig. 1A**). It was previously reported that the H3K4me3:H3K36me3 ratio around H3K4me3 peaks is higher for nucleosomes immediately adjacent to PRDM9 binding sites than for nucleosomes further away,^32^ but this pattern was not apparent when centered on SPO11-oligo hotspots (**Fig. 1A**), suggesting that this is not a robust feature of PRDM9-dependent histone methylation. This pattern was irrespective of whether we applied local normalization.

We previously reported that the apparent symmetry in H3K4me3 disposition around hotspots (see **Fig. 1A**) is an artifact of averaging, and that individual hotspots display a continuum of varying degrees of left–right asymmetry (**Fig. 1B**, left panel).^26^ This feature was also seen for H3K36me3, and the profiles for the two modifications were highly correlated (**Fig. 1B–C**). Hotspots can be classified into three groups on the basis of the spatial relationship between local SPO11-oligo patterns and putative 36-base-pair (bp) PRDM9 binding site (**Fig. 1D**).^26^ H3K36me3 spatial distribution and ChIP signal strength were comparable among the three hotspot classes, again largely indistinguishable from prior findings for H3K4me3 (**Fig. 1D– E**).^26^

Whereas variation in H3K4me3 ChIP signal could account for 40% of the variation in SPO11-oligo counts at hotspots (R^2^ = 0.40),^26^ H3K36me3 could only account for 26% (**Fig. 1F**). This difference is partially due to higher background signal from transcription-dependent H3K36me3: when we eliminated hotpots in genes previously shown to be transcribed in juvenile testes^33^ (where meiotic cells are enriched because of semi-synchronous spermatogenesis), the correlation increased to ∼33% (data not shown). By multiple linear regression, combining data for both histone marks for all hotspots gave a significant but quantitatively small improvement over a model with H3K4me3 alone for predicting SPO11-oligo counts (R^2^ = 0.44; p < 2.2 × 10^-16^, ANOVA; **Table S1**).

PRDM9 has been proposed to trimethylate H3K4 and H3K36 on the same nucleosome at least some of the time.^32^ Taken together, our findings of highly similar H3K4me3 and H3K36me3 patterns (both spatial and quantitative) around SPO11-oligo hotspots agree with this proposal. Conversely, the findings provide little if any support for an alternative hypothesis in which methylation of the two residues contribute to hotspot activity independently.

### Combinatorial effects of ATM and PRDM9 in controlling hotspot heat

Hotspots are just one organizational level among many in the DSB landscape. In the budding yeast *Saccharomyces cerevisiae*, DSB distributions are shaped by multiple factors that work hierarchically and combinatorially.^4,6,34-36^ For example, most yeast hotspots correspond to the nucleosome-depleted regions in gene promoters,^4,37,38^ but how strong a hotspot will be is shaped not only by factors within the hotspot itself, but also by larger-scale chromosome structures operating over distances from tens of kilobases (kb) up to whole chromosomes.^4,36,38-40^ Some of these factors involve feedback circuits that regulate the ability of Spo11 to continue making DSBs depending on whether DSBs have already formed and whether chromosomes are successfully engaging their homologs (reviewed in ref.^41^) The mammalian DSB landscape is thought to be shaped similarly,^24,26,35^ but there has been little if any formal exploration of the degree to which different factors interact with one another. To address this question, we compared ATM- and PRDM9-dependent contributions to the DSB landscape.

The ATM (for <u>a</u>taxia telangiectasia <u>m</u>utated) kinase triggers checkpoint signaling and promotes DSB repair.^42^ Meiotic DSBs activate ATM, which in turn suppresses further DSB formation.^43^ This negative feedback circuit is conserved in *S. cerevisiae*, dependent on the yeast ATM ortholog Tel1.^40,44,45^ In both mouse and yeast, ATM/Tel1-dependent DSB control also shapes the DSB landscape.^26,40^

We hypothesized that ATM-mediated DSB control is independent of PRDM9 activity. We showed previously that in the absence of ATM, nearly all hotspots experience more DSBs but weaker hotspots increase more in heat than stronger ones: the ratio of SPO11-oligo counts in ATM-deficient relative to ATM-proficient samples was negatively correlated with hotspot heat.^26^ By linear regression this correlation accounted for 16% of the variation in SPO11-oligo counts at hotspots found in the C57BL/6J (“B6”) strain (**Fig. 1G)**. Differences between hotspots in their response to *Atm* mutation correlated poorly with their H3K4me3 levels (**Fig. 1H**). Furthermore, a multiple linear regression model that combined measures of PRDM9 activity (H3K4me3 and H3K36me3) with measures of the effects of ATM (*Atm*^*–/–*^:*Atm*^*+/+*^ SPO11-oligo ratio) substantially improved the ability to predict hotspot heat relative to a model incorporating H3 methylation status only (R^2^ = 0.60; **Fig. 1I** and **Table S1**). Taken together, these findings support the idea that ATM and PRDM9 contribute largely independently to determining the heat of individual hotspots. More generally, these findings illustrate the degree to which different factors can interact to shape the DSB landscape.

### Characteristics of DNA sequences around 5′ and 3′ ends of SPO11 oligos

Fine-scale analyses of sequence composition around meiotic DSB sites in budding and fission yeasts revealed non-random base composition but no apparent consensus sequence^4,46-49^. SPO11 generates two-nucleotide (nt) 5′ overhangs,^47,48^ predicting an axis of rotational symmetry at the phosphodiester bond between the first and second position for each mapped SPO11 oligo (**Fig. 2A**).^4^ We examined possible SPO11 biases in mice by orienting and aligning the DNA sequences surrounding each uniquely mapped SPO11 oligo from 9,060 PRDM9-motif-containing hotspots and then evaluating DNA sequence composition around the predicted dyad axis (**Fig. 2A**). Base composition deviated from random at all positions from six nt upstream to 21 nt downstream of the dyad axis (-6 to +21). The bias from +10 to +21 may reflect preferences related to oligo 3′-end formation, discussed further below. The strong bias in the central 12 nt, a region predicted to contact SPO11,^50^ could have reflected preferences of SPO11 itself as inferred in yeasts,^4,49^ but could alternatively reflect signatures of the sequence specificity of PRDM9 (whose binding sites lie very close to many SPO11 cleavage sites (**Fig. 1D**)) or biases of the sequencing and/or mapping methods.

**Figure 2.**
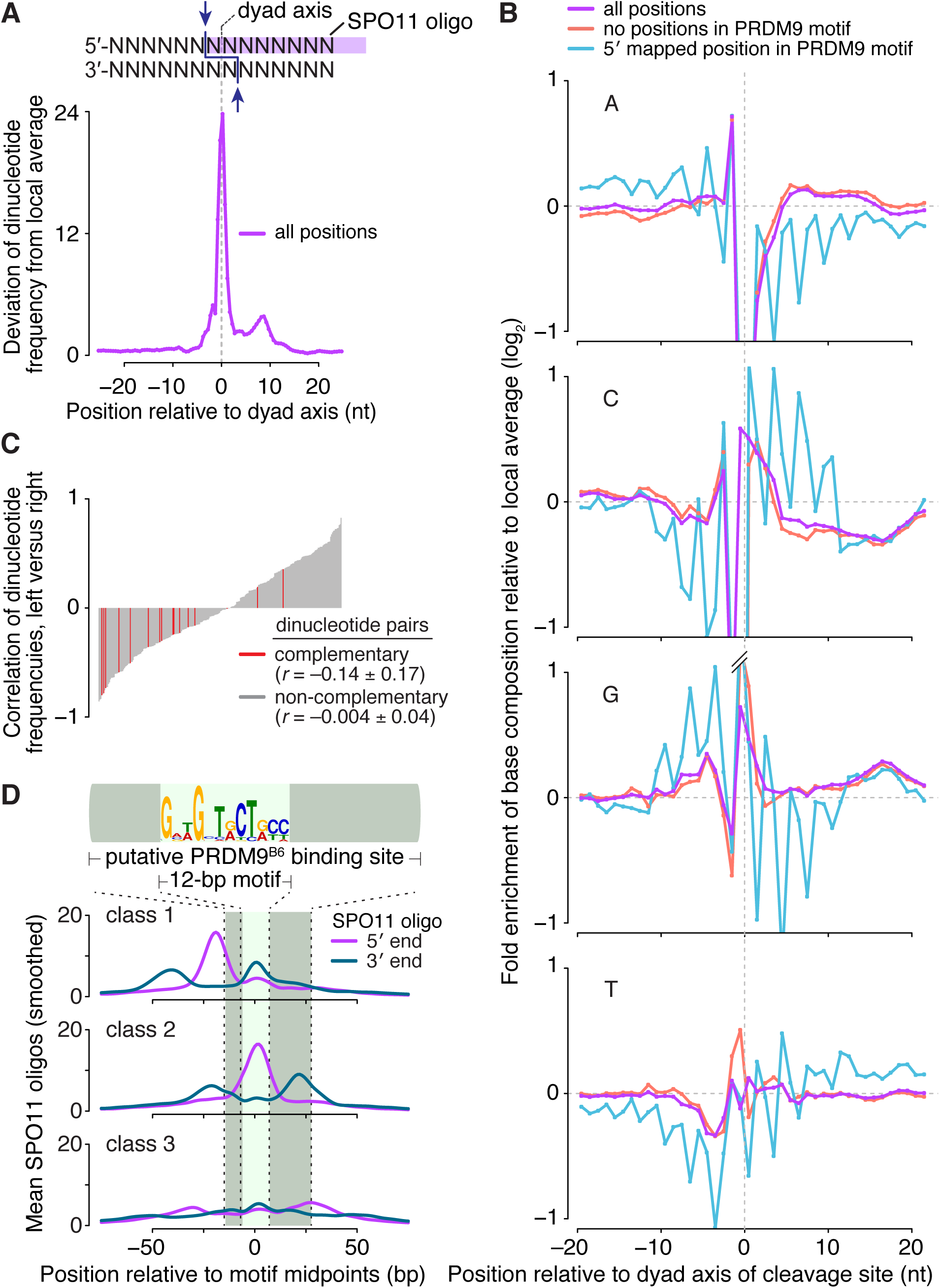
Sequence composition at sites of SPO11-oligo formation and 3′ nicking. (A) (top) Schematic of a SPO11 DSB. Staggered cuts (arrows) by a SPO11 dimer generate two-nucleotide 5′ overhangs, in the middle of which is a two-fold axis of rotational symmetry. (bottom) Non-random dinucleotide composition around SPO11 cleavage sites in 9,060 SPO11-oligo hotspots. At each position, deviation of dinucleotide frequencies from local average was summed. (B) Mononucleotide composition around the 5′ ends of uniquely mapped SPO11 oligos within hotspots. Purple, all cleavage sites; blue, SPO11 oligos with the 5′-mapped end in a PRDM9 motif and without a 5′ C; red, SPO11 oligos without an overlap to a PRDM9 motif and without a 5′ C. Plots are truncated for values outside of the range from −1 to 1. (C) Dinucleotide base composition is not rotationally symmetric around the dyad axis. The dinucleotide composition on the left of cleavage sites was compared to the composition on the right for each of the 256 possible dinucleotide pairs. The correlation coefficients (Pearson’s *r*) were then rank-ordered and displayed as a bar plot with reverse-complementary dinucleotides colored in red. The mean ± SD of the *r* values is shown for reverse-complementary and non-reverse-complementary dinucleotides. (D) The distributions of 3′ ends of SPO11 oligos relative to 5′ ends are similar for the three classes of PRDM9 motifs defined by local SPO11-oligo pattern (see **Fig. 1D**). The 5′ and 3′ ends of SPO11-oligo profiles were smoothed with a 15-bp Hann filter.

We therefore asked if base composition biases near SPO11-oligo mapping positions were rotationally symmetric around the predicted SPO11-dyad axis, as previously demonstrated in *S. cerevisiae*.^4^ However, the base composition was not clearly two-fold rotationally symmetric (**Fig. 2B**). For example, A residues were enriched at the −2 position but there was no reciprocal enrichment of T at +2. We further explored this question by assessing the degree to which the dinucleotide frequencies to the left of the dyad axis correlated with dinucleotide frequencies to the right. To do so, we examined the regions from −3 to −16 and +3 to +16 to avoid the artifactual enrichment or depletion of C-containing dinucleotides around the 5′ end of the mapped SPO11 oligos.^4^ For rotationally symmetric patterns, pairwise comparisons of reverse-complementary dinucleotides should show high, positive values of Pearson’s *r*, as previously observed in *S. cerevisiae*.^4^ However, we found no such enrichment for positive *r* values (**Fig. 2C**), thus the sequence bias around mouse SPO11-oligo sites cannot be clearly ascribed to preferences of SPO11 itself.

To test whether presence of PRDM9 binding motifs was obscuring underlying SPO11-associated base composition bias, we focused on two sets of hotspot-associated SPO11 oligos (after first excluding reads with 5′ ends mapping to C residues to minimize a known technical bias^4^): 1) reads whose 5′ ends mapped within a PRDM9 motif, and 2) reads that did not overlap a PRDM9 motif anywhere along their lengths. Base composition around the two sets displayed distinct patterns in addition to the selected-for depletion of C at the −1 position (**Fig. 2B**). For cleavage sites within PRDM9 motifs there was a pronounced 3-nt periodicity in the frequencies of each mononucleotide (blue lines in **Fig. 2B**). This periodicity presumably reflects the fact that each individual PRDM9 zinc finger recognizes a DNA triplet sequence.^19,20,51^ However, SPO11-oligo sites that did not overlap a PRDM9 motif still displayed little or no evidence of rotationally symmetric base composition (red lines in **Fig. 2B**). Thus, unlike in yeast, we have been unable to discern in mouse a clear signature that we can ascribe to SPO11 preferences for particular base compositions. These findings do not exclude such preferences contributing to cleavage site choice, but if they exist, such contributions appear to be quantitatively weak relative to other sources of non-randomness in fine-scale SPO11-oligo maps.

In *S. cerevisiae*, the 3′ ends of Spo11 oligos are thought to be formed by a combination of endonuclase and 3′→5′ exonuclease activities of the Mre11-Rad50-Xrs2 complex plus Sae2 (MRE11-RAD50-NBS1 and CTIP in mouse).^9,52-55^ Because many SPO11 oligos overlap the positions where PRDM9 binds, we asked whether PRDM9 might influence SPO11-oligo 3′-end formation. On average, oligo 3′ ends were offset by similar lengths from oligo 5′ ends in both directions, reflecting oligos mapping to the top and bottom strands of the DNA (**Fig. 2D**). The offsets of 3′-end positions were as expected for the lengths of mapped SPO11-oligo reads, irrespective of where DSBs occurred relative to PRDM9 binding motifs. This finding implies that PRDM9 has little impact on the nucleolytic processing steps that form the 3′ ends of SPO11 oligos, in turn suggesting that PRDM9 may not be bound to DNA when the MRE11 complex completes its processing function. This inference fits with an earlier proposal that oftentimes PRDM9 has already left its binding sites when SPO11 cleaves the DNA, on the basis of the frequent overlap of SPO11 oligos with PRDM9 binding sites (e.g., **Fig. 1D**).^26^ Recombinant PRDM9 forms long-lived complexes with DNA *in vitro*,^56^ raising the possibility that it is actively displaced from its binding sites *in vivo* by SPO11 and/or other factors.

### DSBs are underrepresented but nonetheless occur frequently within repeated sequences

The mouse genome is replete with repeated sequences.^57,58^ These sequences pose a risk because a meiotic DSB formed within one copy of a repeat has the potential to cause a chromosomal rearrangement by repair with a non-allelic copy. Such non-allelic homologous recombination (NAHR) can lead to deleterious deletions, duplications, and other alterations. Indeed, NAHR-mediated chromosomal rearrangements in the germline are the cause of numerous developmental disorders in humans.^3,59-61^ In budding yeast, DSB formation tends to be repressed within repetitive DNA genome-wide,^4,62^ perhaps as a mechanism to protect against NAHR. We hypothesized that this strategy is also found in mice.

We asked whether SPO11 oligos were less likely to arise from repeated sequences than expected if DSBs were formed in proportion to the relative amount of genomic space occupied by repeats. SPO11 oligos were assigned to two sequence classes: “repeat” and “non-repeat” (**Fig. 3A**). The repeat class, totaling 1,286,319,054 bp, encompassed two sub-groups: interspersed repeats such as transposable elements and low complexity sequences compiled by RepeatMasker (www.repeatmasker.org), which together comprise 45% of the genome;^57^ and segmental duplications, defined as genomic segments sharing ≥90% sequence identity over ≥1 kb, which collectively occupy 8.1% of the genome.^58^ These two sub-groups are partially overlapping (4.7% of the genome). The non-repeat class encompasses the remainder of the genome and totals 1,366,448,147 bp.

**Figure 3.**
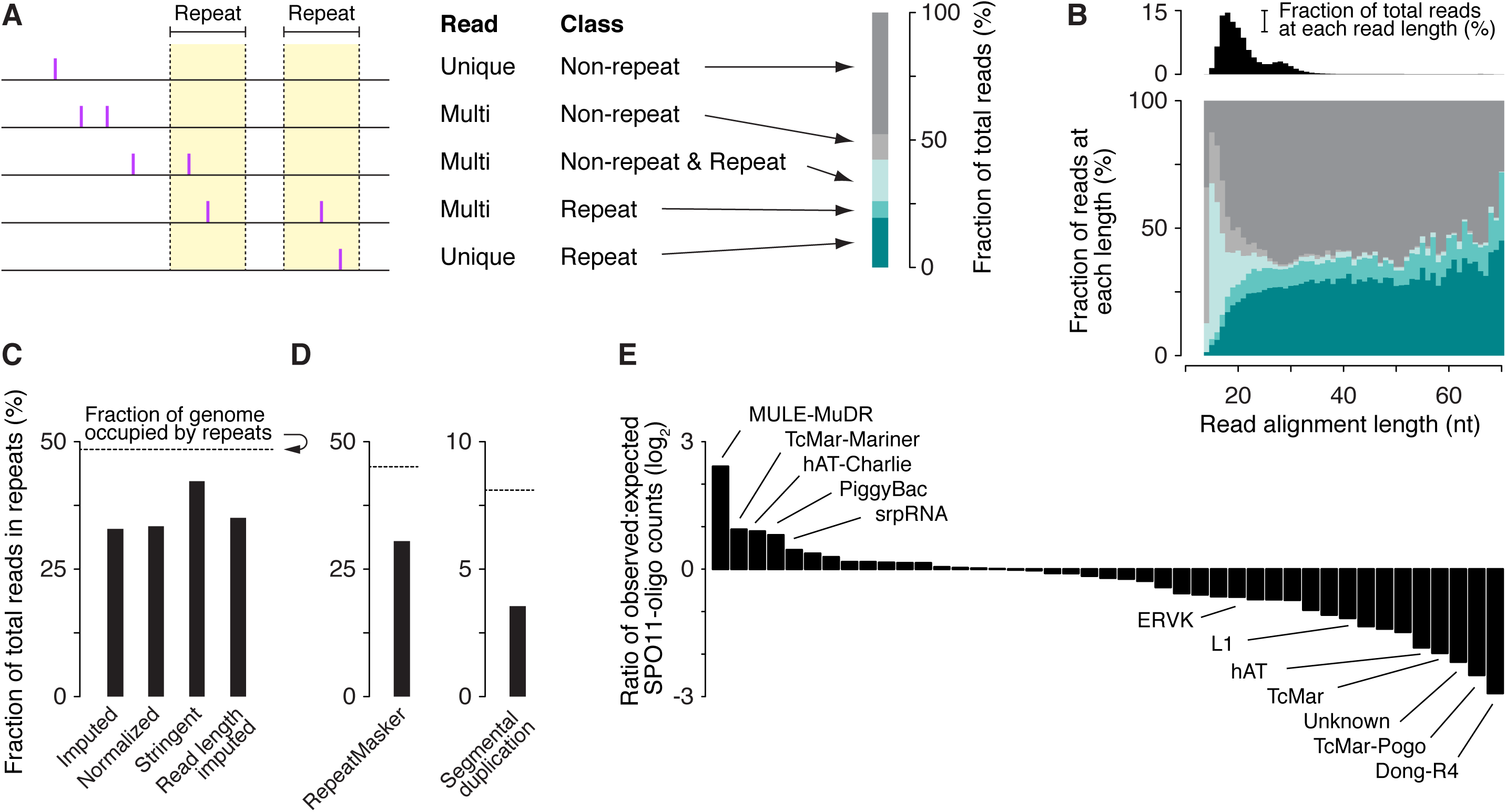
DSBs in repeated sequences. (A) Five categories of SPO11-oligo mappability. 67.2% of SPO11 oligos could be assigned unambiguously to a unique location in the reference genome within either the non-repeat class (dark gray) or the repeat class (dark turquoise). The remaining 32.8% mapped to multiple places in the reference genome; half of these could be placed in either the non-repeat class (i.e., all mapped positions are in non-repeat sequences; light gray) or the repeat class (i.e., all mapped positions are in the repeat class; turquoise). However, the remaining half of the multi-mappers (16.3% of all mapped reads) could have derived from either class because reads mapped to both repeated and non-repeated sequences (light turquoise). (B) (top) Histogram of SPO11-oligo read lengths (adapted from ref ^26^ with permission). (bottom) Percentages of the five categories as a function of read length. Short reads (< 20 nt) were highly enriched for multi-mapped reads, suggesting that many such SPO11 oligos were mapped to multiple places solely because of their short length. (C) Percentages of SPO11 oligos mapped to repeat sequences, estimated by four methods. Imputed: Reads were assigned in proportion to the number of unique reads in the neighboring area (method described in ref.^26^). Normalized: Each multi-mapped read was divided evenly among its mapped positions. Stringent: All multi-mapped reads that could have derived from either repeat or non-repeat sequences were assigned entirely to that respective repeat class. This approach likely overestimates the DSB frequency in repeat sequences and therefore provides the most conservative estimate of the degree to which DSB formation is suppressed in repeats. Imputed read length: Because the mappability of short reads is less certain (B), we included from the imputed map only reads with lengths of 25–50 nt and recalculated the fraction of imputed reads in repeated sequences. This method represents the highest confidence estimate of the relative burden of DSBs in sequences annotated as repeats. (D) DSB frequencies in two sub-groups of repeat sequences. SPO11-oligo frequencies were lower than expected in interspersed repeats and low complexity DNA sequences (“RepeatMasker”) and in segmental duplications. (E) DSB frequencies in families of repeat sequences annotated in RepeatMasker. Overlapping repeats of the same family were merged before calculation. For each repeat family, SPO11-oligo counts per base pair were summed and their enrichment was calculated relative to expected values (see Materials and methods). The most extreme examples are labeled, as are L1 and ERVK families, which include elements that are putative DNMT3L targets.^63^ The identities, calculated enrichment values, and fractions of the genome occupied for all of the families analyzed are provided in **Table S2**.

Reads that map to multiple locations are conventionally excluded from Next Generation Sequencing studies because they cannot be assigned unambiguously. However, the exclusion of multi-mapped reads in this analysis would result in an underestimation of the DSB frequency in repeated sequences. Some SPO11 oligos can be mapped uniquely even to repeat-class sequences (**Fig. 3A**), because individual copies of repetitive elements can often be differentiated from one another by DNA sequence variants. Conversely, SPO11-oligos can be assigned to multiple locations that lie within non-repeat sequences as a consequence of the short length of SPO11-oligos and the low complexity of the mouse genome relative to a truly random DNA sequence of the same size (**Fig. 3A–B**).^26^ Thus, multi-mapped reads that map to both repeat and non-repeat sequences cannot be unambiguously placed into either class.

We therefore used four methods with varying stringency to allocate multi-mapped reads among the repeat and non-repeat classes (**Fig. 3A–C**). By all four approaches, the frequency of SPO11-oligos in repeat sequences was lower than expected based on the fraction of the genome occupied by such sequences, and this was true for both sub-groups of the repeat class (**Fig. 3A–D**). This apparent DSB suppression was the case for many families of interspersed repeats, including retrotransposons such as the L1 (LINE1) elements L1Md_A, L1Md_Gf and L1Md_T, and the ERVK family LTR (long terminal repeat) elements IAPEz and MMERVK10C, whose meiotic transcription is suppressed by DNA methylation targeted to them by DNMT3L, a catalytically inactive member of the Dnmt3 DNA methyltransferase family (**Fig. 3E** and **Table S2**).^63^ Interestingly, however, several repeat element classes had higher SPO11-oligo frequencies than expected, including sequences that RepeatMasker annotated as homologous to the DNA transposon subfamilies MULE-MuDR, TcMar-Mariner, hAT-Charlie, and PiggyBac.

Two main conclusions arise from this analysis. The “glass half empty” viewpoint is that, on average, repetitive DNA sequences incur fewer DSBs than expected by chance. The “glass half full” viewpoint is that this apparent DSB suppression, while it may be strong for individual repeats, is quantitatively modest in global terms: assuming 200–300 DSBs per spermatocyte,^64^ as many as 65–100 DSBs are formed in each meiosis within sequences that have at least some repetitive character. The extent to which meiotic NAHR depends on the percent sequence similarity and length between repeats remains unknown. Thus, for many or most of these DSBs, it is likely that sequence differences from non-allelic copies prevent NAHR from occurring. However, it is likely that at least some of these DSBs pose significant risk for NAHR in every meiosis. The mechanisms that would allow a cell to tolerate this risk remain poorly understood.^3,60,65,66^

It appears that active mechanisms may disfavor DSB formation within specific repeat families, e.g., as a result of targeted DNA methylation and heterochromatinization.^63^ However, because SPO11-oligo underrepresentation appears to be broadly true of many repeat classes, we propose that this feature of the DSB landscape reflects in part a selective constraint against having too many recombination events involving repeats. More specifically, we hypothesize that PRDM9 proteins that target repetitive DNA too robustly may not be compatible with fertility, thus only those *Prdm9* alleles that do not confer too high a burden of repeat-associated DSBs are likely to be found in populations.

A corollary of this hypothesis is that inappropriately high DSB levels in repeats may contribute to the infertility seen in some mouse hybrid strains, which is caused by incompatibility between the *Prdm9* allele of one strain and the genome of the other.^24,67-69^ Prior studies have established that small differences in total DSB numbers can spell the difference between successful meiosis and infertility caused by spermatogenic failure.^64,70^ These findings have been interpreted to mean that there is a threshold in the number of recombination events needed to support successful chromosome pairing, synapsis, and segregation in mice.^64,70-72^ DSBs within repeats may not contribute effectively to chromosome pairing and may in fact interfere with pairing by stabilizing illegitimate interactions between non-homologous chromosomes. Thus, given the surprisingly high apparent fraction of DSBs in repeats in the B6 strain (**Fig. 3**), it may take only a modest further increase in this burden to trigger catastrophic meiotic defects.

Nonetheless, it is noteworthy that PRDM9 binding sites are enriched in certain classes of repeated sequence, for example in human L2 LINEs, AluY elements, and the retrovirus-like retrotransposons THE1A and THE1B.^20,73^ Thus, paradoxically, it may be that PRDM9 functions include the preferential targeting of certain repeats, perhaps as part of the cell’s attempts to limit the spread of certain selfish genetic elements. If so, this function must be balanced against formation of too many DSBs in repeats. We speculate that this balance is an important component of the remarkable evolutionary dynamics of PRDM9, including multiple independent apparent losses of *Prdm9* genes in a number of vertebrate lineages.^74-78^

### Histone H3 methylation and the presence of ssDNA

Single-stranded DNA sequencing (SSDS) maps meiotic DSBs by immunopurification and deep-sequencing of DMC1-bound ssDNA from spermatocytes.^21^ Because SSDS captures an intermediate that arises after resection, SSDS and SPO11-oligo data can be combined to explore resection.^26^ Nucleosomes can impede resection by exonucleases *in vitro*,^79^ and chromatin structure strongly influences meiotic resection patterns *in vivo* in *S. cerevisiae*.^13^ Given the pronounced asymmetry in PRDM9-dependent histone H3 methylation at many hotspots (**Fig. 1B**), we asked if this feature of the local chromatin structure was correlated with resection patterns. Interestingly, at asymmetrically methylated hotspots, SSDS signal strength was 8% higher on average on the sides of the hotspots with highest H3K4me3 (and H3K36me3) ChIP signal (**Fig. 4A–C**). The distance over which the SSDS signal spread away from hotspots was not apparently different, however, thus the lengths of resection tracts are not correlated with H3 methylation status. [Note that the H3 methylation ChIP signal does not allow conclusions to be drawn about total nucleosome occupancy (methylated plus unmethylated).]

**Figure 4.**
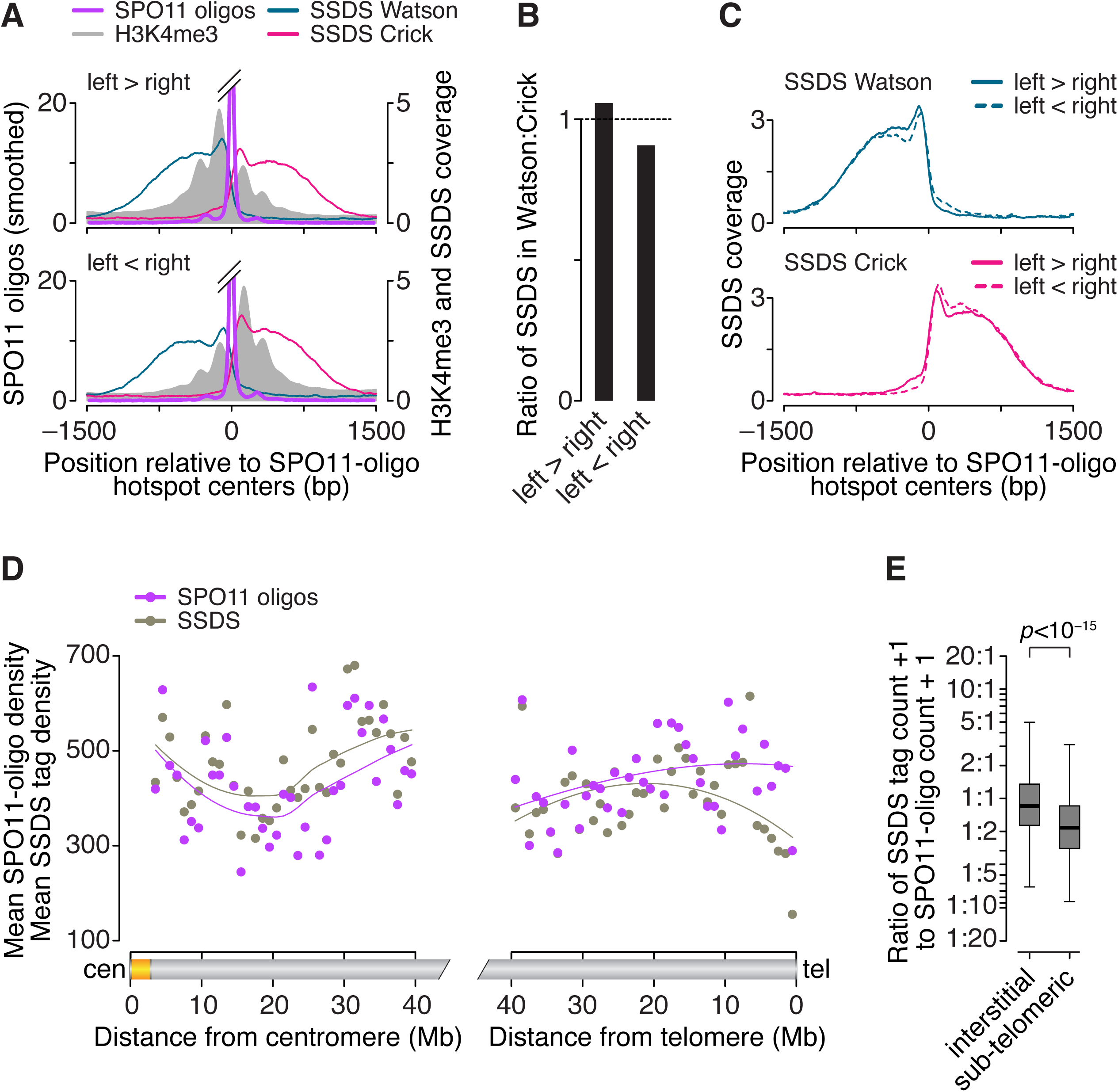
Factors influencing DSB processing. (A) Spatial correlation of SSDS coverage (data from ref. ^21^) with methylated nucleosomes. Each panel shows the mean of locally normalized profiles across the 20% of hotspots with the most asymmetric H3K4me3 patterns (left > right in top panel; right > left in bottom panel). The same local normalization factor was applied to Watson and Crick SSDS reads for each hotspot so that signal strengths on Watson and Crick were comparable. (B) Ratio of sum of locally normalized SSDS coverage in 3,001-bp windows around the centers of the subsets of hotspots from panel A. (C) SSDS coverage was concordant with H3K4me3 signal, albeit weakly. Each panel shows the mean of locally normalized profiles on Watson (top) and Crick (bottom) strands in the hotspots subsetted by H3K4me3 asymmetry. Local normalization was applied separately to Watson and Crick reads so that signal strengths were comparable between the subsets. (D) SPO11 oligos and SSDS coverage display similar density patterns in centromere-proximal regions (left) but not in centromere-distal regions (right). Points are densities of SPO11 oligos (reads per million (RPM) per Mb) and SSDS tag counts (tags per million (TPM) per Mb), within coordinates defined by SSDS hotspots, in 1-Mb windows, averaged across all 19 autosomes. In the right panel, one SPO11-oligo outlier is not shown. (E) Ratios of SSDS tag counts to SPO11-oligo read counts in SSDS hotspots differ between by autosomal sub-chromosomal domain. “Sub-telomeric” is defined as the centromere-distal 5 Mb of each autosome. “Interstitial” is all other autosomal regions. One tag count and one SPO11-oligo read count were added to each hotspot. Boxplot outliers are not shown. P value is from Wilcoxon rank sum test.

SSDS signal strength reflects both the number of DSBs in the population and the lifespan of DMC1-bound ssDNA.^26^ SSDS signal on the left side of hotspots derives from precisely the same DSBs as the SSDS signal on the right, therefore the observed left-versus-right asymmetry in SSDS signal cannot be ascribed to differences in DSB frequency or timing of DSB formation. Instead, the asymmetry may indicate that DMC1 remains bound for a shorter time on average on the low-methylation side of hotspots. For example, this side might tend to engage in strand exchange and DMC1 dissocation earlier than the high-methylation side. Effects on recombination efficiency have been inferred for PRDM9-dependent histone methylation of the intact recombination partner.^24^ Alternatively, the asymmetry might reflect systematic differences in the amount of DMC1 bound to the ssDNA on the two sides. We also cannot exclude the possibility that the asymmetry derives from technical biases that affect coverage maps in both SSDS and H3 methylation ChIP experiments.

### Non-centromeric ends of autosomes display unusually low SSDS:SPO11-oligo ratios

Previously, we found that hotspots on sex chromosomes show a markedly higher ratio of SSDS to SPO11-oligo counts than hotspots on autosomes.^26^ SPO11 oligos have a long lifespan relative to the length of prophase,^43^ so (unlike SSDS) SPO11-oligo mapping is thought to be relatively insensitive to variation in timing of DSB formation or lifespan of DSBs. Thus, it was inferred that DSBs have a longer average lifespan on the non-homologous parts of the sex chromosomes than on autosomes, presumably because of delayed repair caused by absence of a homologous chromosome plus a temporary barrier to using the sister chromatid as a recombination partner.^23,26^

Following on this precedent, we extended the comparative analysis of SPO11-oligo and SSDS maps to large domains of autosomes. Mouse chromosomes are acrocentric, i.e., their centromeres lie close to one end of each chromosome. On average, the centromere-proximal 40 megabases (Mb) of autosomes showed similar patterns in both mapping methods, that is, the ratio of SSDS signal to SPO11-oligo counts was indistinguishable from genome average (**Fig. 4D**). (Note that this analysis excludes the large blocks of repetitive satellite DNA that make up the pericentric heterochromatin. These regions are not readily accessible to deep-sequencing methods, but are also known from cytological data to experience few if any meiotic DSBs.^80^) In contrast, the 5 Mb adjacent to centromere-distal telomeres displayed lower mean SSDS coverage than expected from the SPO11-oligo density (**Fig. 4D–E**). This finding suggests that there are systematic differences in the formation, processing, and/or repair of DSBs in these sub-telomeric zones relative to interstitial regions. This pattern could be explained if DSBs tend to form later near telomeres than elsewhere, or if they form with similar timing but with shorter resection distance or less DMC1 loading. Alternatively, sub-telomeric DSBs may tend to be repaired more quickly, thereby reducing the lifespan of DMC1-bound ssDNA. Different repair kinetics could arise from regional differences in the physical proximity of homologs, for example, via tethering to the nuclear envelope.^81^

### Reciprocal crossover asymmetry and polymorphisms in the PRDM9 binding motif

Both crossover and noncrossover recombination outcomes can be accompanied by gene conversion of allelic differences around the DSB site (**Fig. 5A**). For crossovers, presence of a gene conversion tract causes the breakpoints on the two recombinant chromosomes to lie at different positions, flanking the conversion tract (**Fig. 5A**). In mice, crossover breakpoints at several recombination hotspots have been characterized by fine-scale analyses of recombinant products in sperm DNA isolated from F_1_ hybrids of B6 with other strains.^82-88^ In these studies, breakpoints were assayed after PCR amplification of recombinant molecules using allele-specific primers in each of the two orientations (i.e., forward primers specific for B6 polymorphisms with reverse primers specific for the other strain; and vice versa). If DSB formation is about equally likely on the two homologous chromosomes, then the distribution of crossover breakpoints will be similar for both PCR orientations. But if DSB formation is biased in favor of one allele, then the breakpoint distribution detected in one orientation will be shifted relative to breakpoints amplified in the other orientation (**Fig. 5A**).^84,89,90^ Such reciprocal crossover asymmetry was observed at the *A3* hotspot in B6×DBA/2J (“DBA”) hybrids and was explained by a sequence polymorphism that resulted in differences in the binding of PRDM9 to the B6 and DBA alleles at *A3*.^84,91^

**Figure 5.**
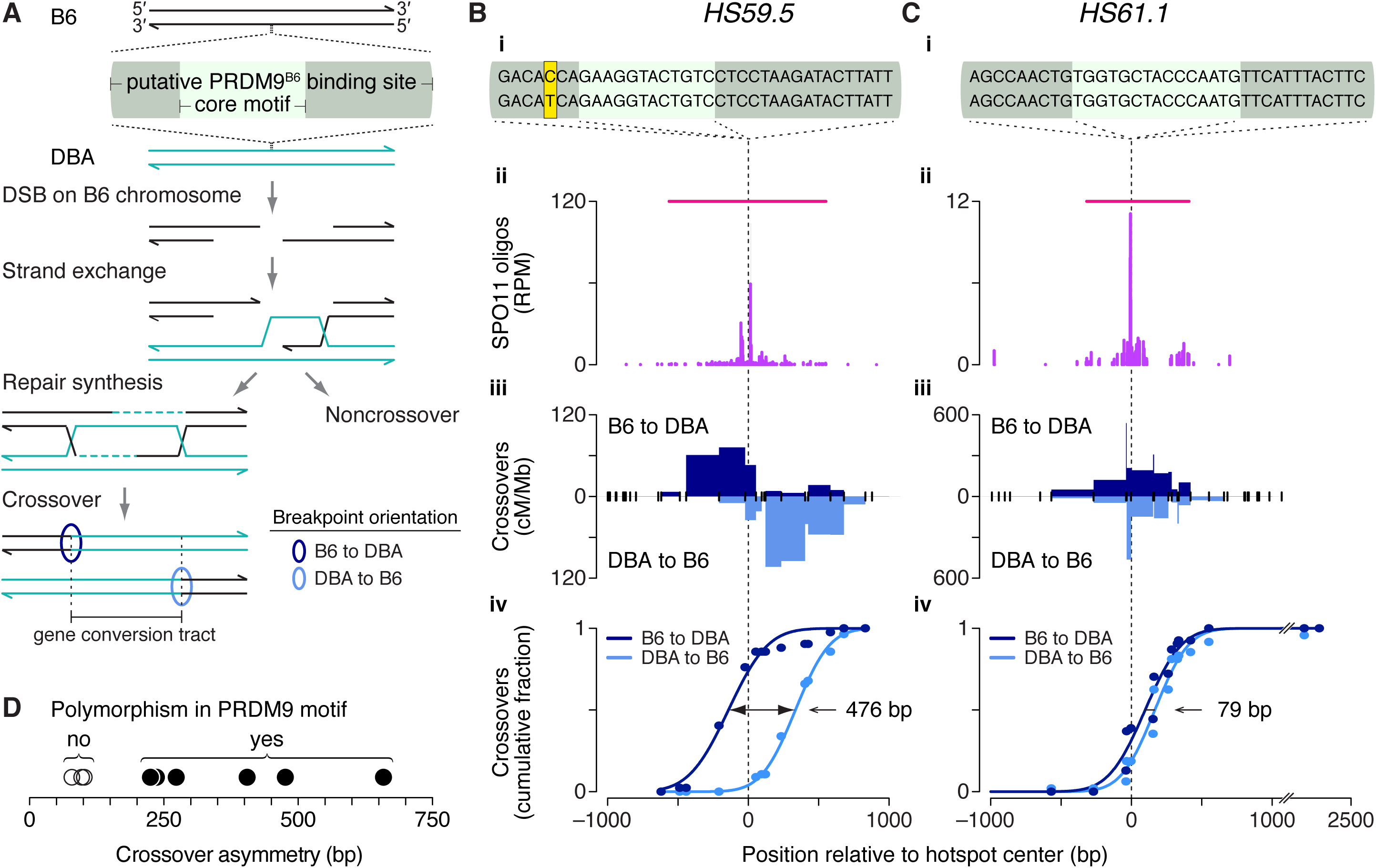
Crossover asymmetry in hotspots with polymorphisms in putative PRDM9 binding sites. (A) Model for crossover asymmetry. A sequence polymorphism in a PRDM9 binding site may affect relative DSB frequencies on the two hotspot alleles, manifested as asymmetry in the locations of crossover breakpoints. If DSBs are preferentially formed on the B6 chromosome of a B6×DBA hybrid mouse, crossover breakpoints will tend to lie to the left of the hotspot center when recombinant products are assayed after PCR amplification in the B6-to-DBA orientation, and will tend to lie to the right when amplified in the DBA-to-B6 orientation. (B–C) Examples of crossover hotspots with (B) or without (C) crossover asymmetry. (i) B6 (top) and DBA (bottom) sequences of putative 36-bp binding sites for PRDM9^B6^ at hotspot centers. The nucleotides shaded in yellow in *HS59.5* highlight a polymorphism between the B6 and DBA haplotypes. In *HS61.1*, the PRDM9 motif shown is on the Crick strand. (ii) SPO11-oligo maps. Red lines indicate SPO11-oligo hotspots. (iii) Crossover breakpoints (densities expressed as centiMorgans (cM) per Mb) mapped by allele-specific PCR on sperm DNA in the B6-to-DBA (top) and DBA-to-B6 (bottom) orientation.^86,87^ Ticks represent tested polymorphisms. (iv) Cumulative distributions of crossover breakpoints with fitted Gaussian curves. The number indicates the distance between the two curves at the midpoint for each cumulative plot. Vertical dashed lines indicate hotspot centers. For hotspot *HS61.1*, zero values in both orientations at outlier position −1130 bp are not shown. (D) Crossover asymmetry is associated with presence of polymorphisms in putative PRDM9 binding sites at hotspots (**Table S3**). Crossover asymmetry was defined for each locus as the absolute difference between the midpoints of cumulative crossover breakpoint maps in the two orientations.

We asked if polymorphisms within PRDM9 binding sites could explain crossover asymmetries observed at other hotspots. Of the 15 published crossover hotspots we examined previously,^26^ 13 contained at least one putative PRDM9 binding motif in B6 within 250 bp of the center of a matched SPO11-oligo hotspot. In all 13 cases, a motif was similarly identified in the allelic sequence in the other strain of the tested F_1_ hybrid (DBA or A/J). Four of these crossover hotspots were excluded from further analysis: three for which crossovers had been assessed in only one orientation and one that encompassed two prominent SPO11-oligo clusters. For the remaining nine crossover hotspots, we examined putative PRDM9 binding sites — defined as the 36 bp encompassing the hotspot-enriched motif^26^ — for polymorphisms between B6 and DBA or A/J (**Fig. 5B–D**).

At *HS59.5*, where the skew in the distribution of crossover breakpoints was 476 bp (**Fig. 5Biii–Biv**), we observed a B6–DBA sequence polymorphism in the putative PRDM9 binding site (**Fig. 5Bi**). In contrast, *HS61.1* displayed no B6–DBA sequence differences in the PRDM9 binding site, and showed essentially no crossover asymmetry (**Fig. 5C**). This pattern extended to all hotspots examined: crossover asymmetry was minimal for three hotspots where the two strains had identical putative PRDM9 binding sites, but each of the six strains with a polymorphism(s) showed reciprocal crossover asymmetry ranging from 225 to 659 bp (**Fig. 5D** and **Table S3**). We infer that the skew in the distribution of crossover breakpoints reflects differential binding of PRDM9 to, and histone trimethylation of, each pair of alleles. *In vitro* assays would be useful for comparing the binding efficiencies of PRDM9 to each allele. We note that in many cases the only polymorphisms present are outside the most conserved nucleotides of the core motif, consistent with non-core positions contributing to DNA binding affinity *in vivo*, as has been shown *in vitro*.^92^

## Materials and methods

### Datasets

We used SPO11-oligo, SSDS and H3K4me3 data from GEO accession numbers GSE84689, GSE35498 and GSE52628, respectively.^21,22,26^ Pre-processing of these data is described elsewhere.^26^ H3K36me3 and RNA-seq data were downloaded from GSE76416 and GSE61613, respectively,^32,33^ and were converted to mouse genome assembly GRCm38/mm10 sequence coordinates using the UCSC Genome Browser LiftOver tool (http://genome.ucsc.edu/cgi-bin/hgLiftOver).

### Quantification and statistical analyses

Statistical analyses were performed using R versions 3.2.3 to 3.3.1 (http://www.r-project.org). Statistical parameters and tests are reported in the figures and legends. In cases where outliers were removed for plotting purposes, none of the data points were removed from the statistical analysis. For crossover breakpoint analyses, cumulative breakpoint distributions with fitted Gaussian curves were determined using GraphPad Prism version 7.

### Base composition analysis

To examine base composition at 5′ ends of SPO11 oligos, we used all uniquely mapped reads from the wild-type B6 dataset. The same conclusions were reached when we used only reads without a 5′-C ambiguity. To compare base composition within and outside of PRDM9 binding sites, we focused on the 201 bp around the peaks of 9,060 SPO11-oligo hotspots with an identified primary PRDM9 motif.^26^ To circumvent any contribution from the 12-bp PRDM9 core motif sequence, we eliminated all reads that overlapped part or all of the 9,865 instances of the motifs (some hotspots had more than one motif). Dinucleotide frequency and correlations for two-fold rotational symmetry were calculated as reported previously.^4^

### Repetitive sequence analysis

Interspersed repeat sequences such as transposons and simple sequence repeats were as annotated by RepeatMasker. Segmental duplications were as defined by Eichler and colleagues.^58^ Tables of RepeatMasker repeats and segmental duplications for mouse genome assembly GRCm38/mm10 were downloaded from the UCSC Table Browser (http://genome.ucsc.edu/cgi-bin/hgTables). To estimate expected SPO11-oligo counts for **Fig. 3E**, we shuffled repeat locations one million times within the mappable portion of the genome and tallied the SPO11 oligos that fell within the shuffled positions. Because of the substantial difference in overall SPO11-oligo densities between autosomes and sex chromosomes,^26^ autosomal repeats were shuffled among autosomes and sex-chromosome repeats were shuffled within their chromosome of origin. We excluded unplaced and unassigned contigs from the analyses. Note that RepeatMasker includes annotations for “rRNA” sequences, but the ribosomal DNA repeat is not assembled in the reference genome (GRCm38/mm10). We excluded these elements from the analysis of specific interspersed repeat element families in **Fig. 3E** but included them for the global analyses in **Fig. 3A–D**.

### PRDM9 binding motifs in crossover hotspots

To identify instances of the previously published^26^ primary or secondary PRDM9 motif around 15 crossover hotspots, we used MAST^93^ to query 501-bp sequences in B6 and A/J or DBA around the centers of matched SPO11-oligo hotspots (**Table S3**). For crossover hotspot *HS18.2*, which does not have a matched SPO11-oligo hotspot, we queried the 501-bp sequence around the midpoint of the region encompassing all crossover breakpoints. This query identified primary and/or secondary PRDM9 motifs in 13 of the 15 crossover hotspots.

To assess crossover asymmetry in hotspots with PRDM9 motifs, we used nine informative loci for which crossover data were available in both orientations and SPO11 oligos were clustered to one zone. The nine crossover hotspots were on chromosome 1 (*A3*, central and distal) and chromosome 19 (*HS18.2*, *HS22*, *HS23.9*, *HS59.5*, *HS61.1* and *HS61.2*). At each of the nine hotspots, we re-mapped previously published crossover data^82,84,86-88^ to B6 coordinates (GRCm38/mm10), then plotted cumulative distributions of crossover breakpoints in each orientation with fitted Gaussian curves. The skew at each hotspot was determined by the distance between the two curves at the midpoint for each cumulative plot.

#### Abbreviations

DSB: double-strand break
ssDNA: single-stranded DNA
ChIP: chromatin immunoprecipitation
NAHR: non-allelic homologous recombination
SSDS: single-stranded DNA sequencing

## Acknowledgments

We thank members of the Keeney and Jasin labs for helpful discussions.

---

**Supplemental Table 1.**
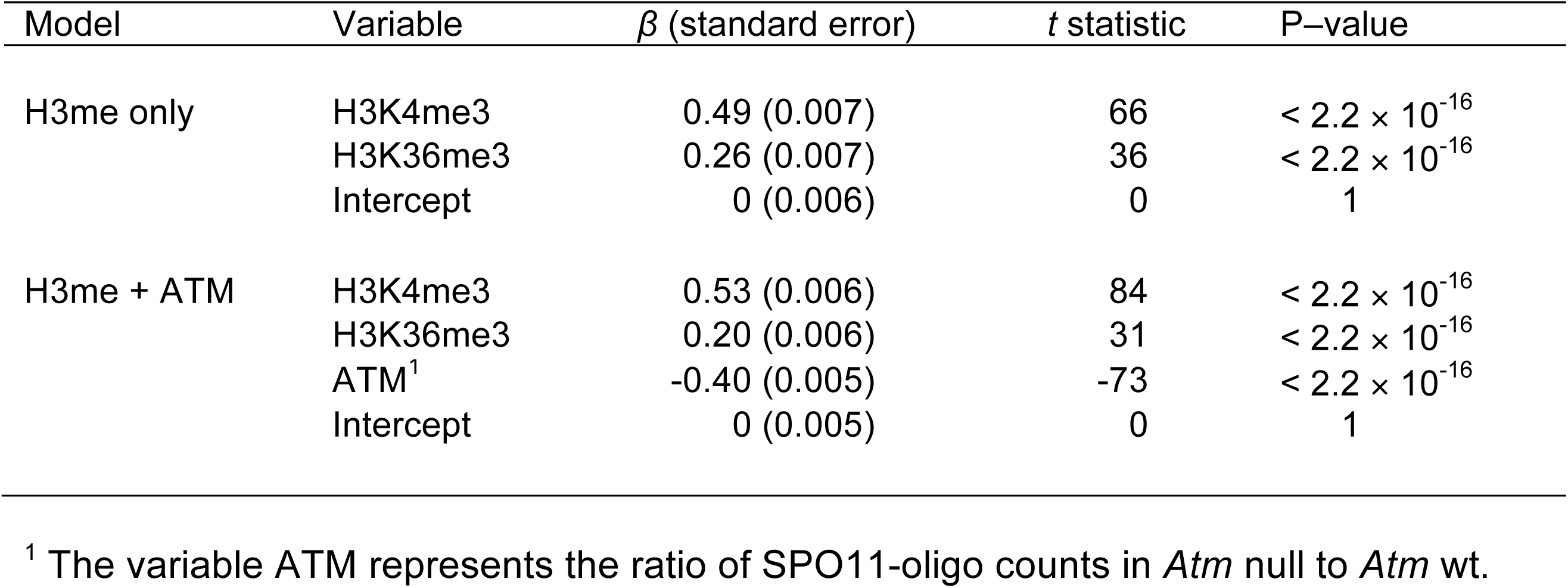
Multiple regression results. Multiple linear regression was performed with log-fold change in SPO11-oligo density around hotspot centers as the dependent variable and the indicated features as the independent variables. SPO11-oligo counts in B6, *Atm* null and *Atm* wt, H3K4me3 tag counts, and H3K36me3 coverage were summed in 1,001-bp windows around B6 hotspot centers. One count was added to each hotspot in *Atm* null and *Atm* wt SPO11-oligo counts, H3K4me3 tag counts, and H3K36me3 coverage counts. The regression analyses were performed on the log-transformed data. Estimates of the standardized regression coefficients (*β*) are shown, along with *t* statistics and P–values based on the standardized coefficients. For the “H3me only” model (H3K4me3 and H3K36me3), multiple R^2^ = 0.4, adjusted R^2^ = 0.4, F = 5,430 on 2 and 13,957 degrees of freedom, and P < 2.2 × 10^-16^. For the “H3me + ATM” model, multiple R^2^ = 0.6, adjusted R^2^ = 0.6, F = 6,748 on 3 and 13,956 degrees of freedom, and P < 2.2 × 10^-16^. Both models were different from the “H3K4me3 only” model (ANOVA P–value < 2.2 × 10^-16^ for each of the two models). Their AIC (akaike information criterion, a measure of a quality of models) values are 47,331 and 42,885, respectively, indicating that the ATM factor improved the fit of the model.

| Model | Variable | $\beta$ (standard error) | *t* statistic | P-value |
| --- | --- | --- | --- | --- |
| H3me only | H3K4me3 | 0.49 (0.007) | 66 | $< 2.2 \times 10^{-16}$ |
| | H3K36me3 | 0.26 (0.007) | 36 | $< 2.2 \times 10^{-16}$ |
|  | Intercept | 0 (0.006) | 0 | 1 |
| H3me + ATM | H3K4me3 | 0.53 (0.006) | 84 | $< 2.2 \times 10^{-16}$ |
| | H3K36me3 | 0.20 (0.006) | 31 | $< 2.2 \times 10^{-16}$ |
| | ATM<sup>1</sup> | -0.40 (0.005) | -73 | $< 2.2 \times 10^{-16}$ |
|  | Intercept | 0 (0.005) | 0 | 1 |

<sup>1</sup> The variable ATM represents the ratio of SPO11-oligo counts in *Atm* null to *Atm* wt.

**Supplemental Table 2.**
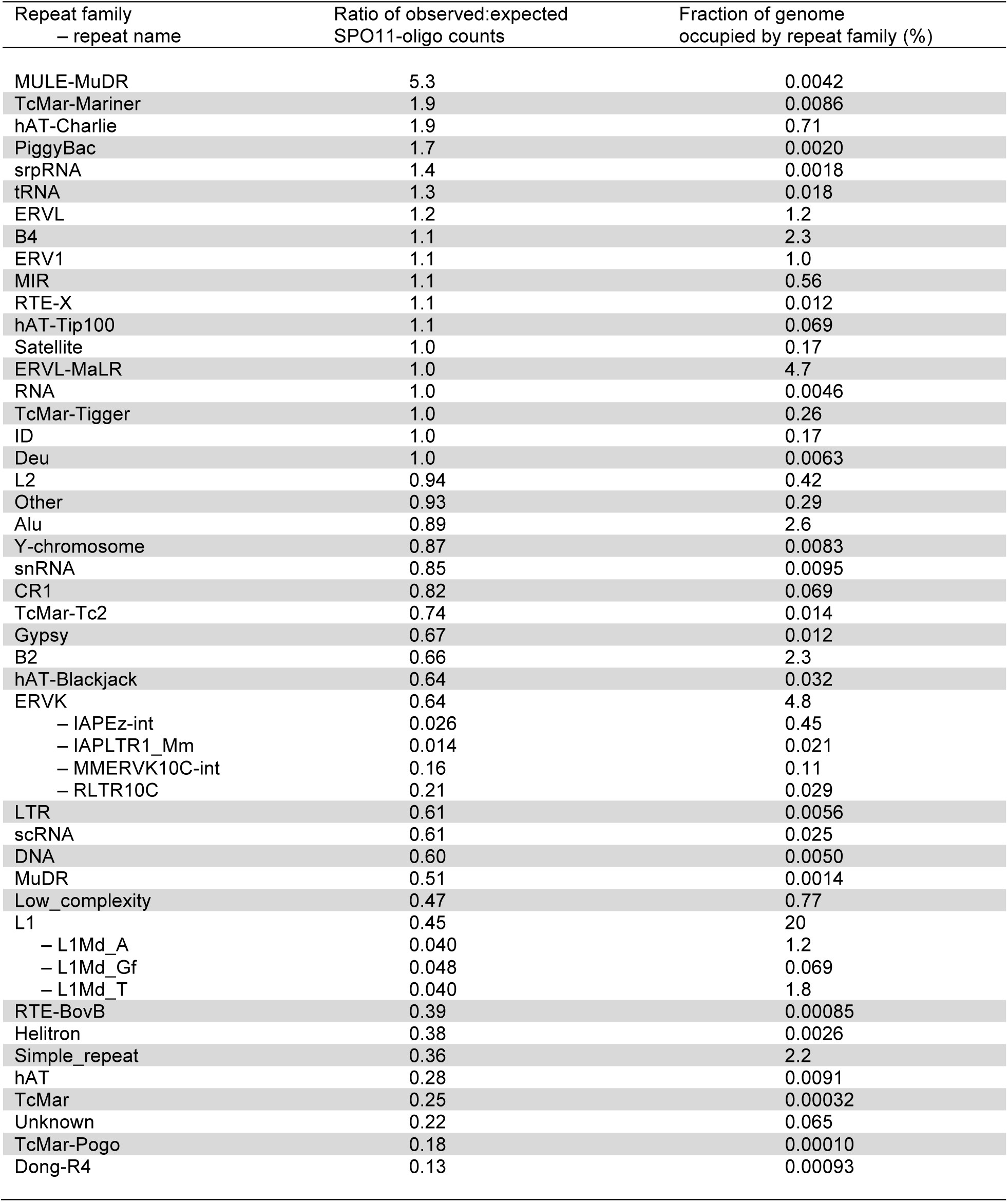
DSB frequencies in interspersed repeats. For each repeat annotated in RepeatMasker, SPO11-oligo counts per base pair were summed and their enrichment was calculated relative to expected values (see Materials and methods). ERVK and L1 repeat family elements whose meotic transcription is suppressed by DNMT3L^63^ are indicated.

**Supplemental Table 3.**
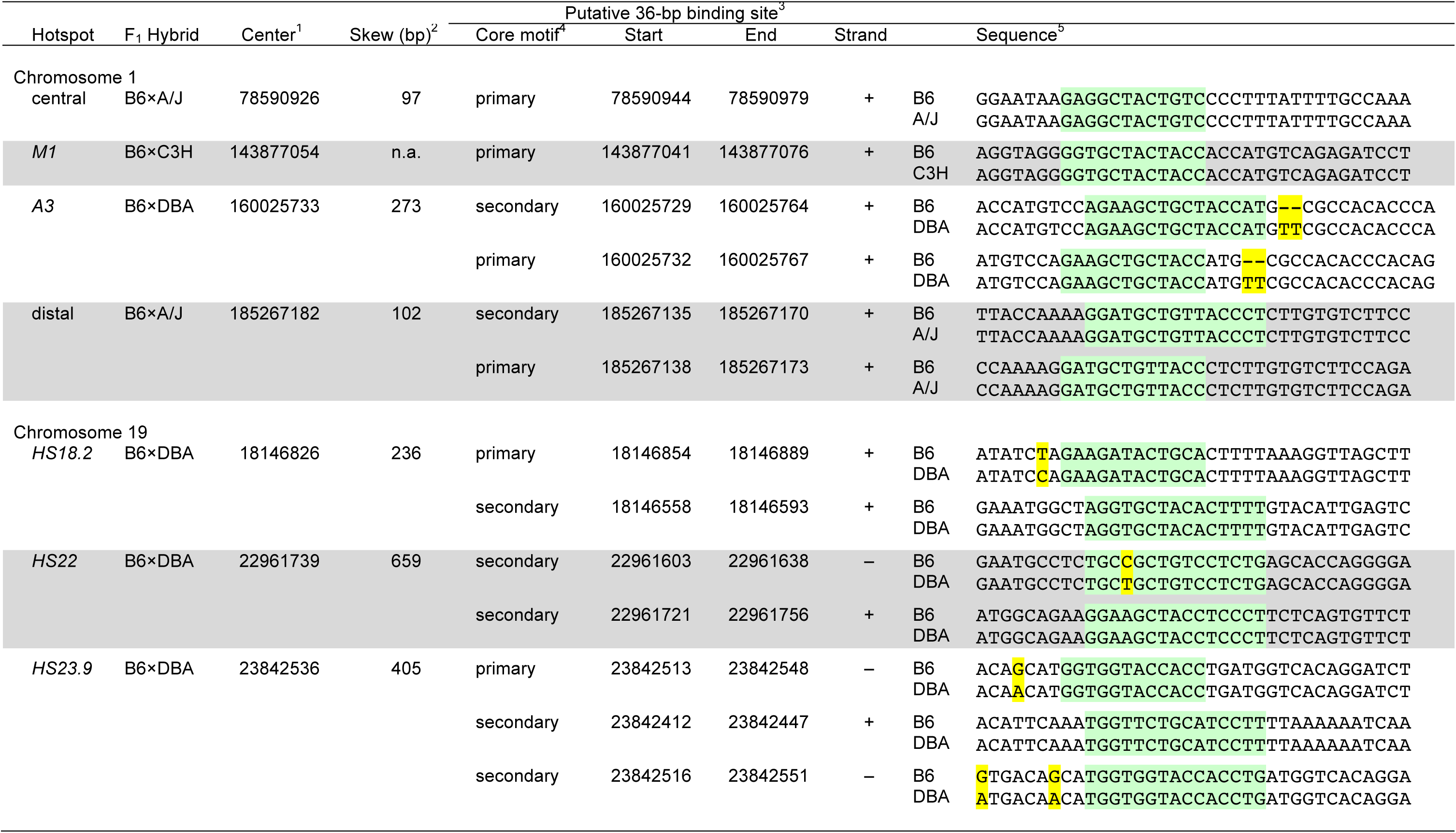
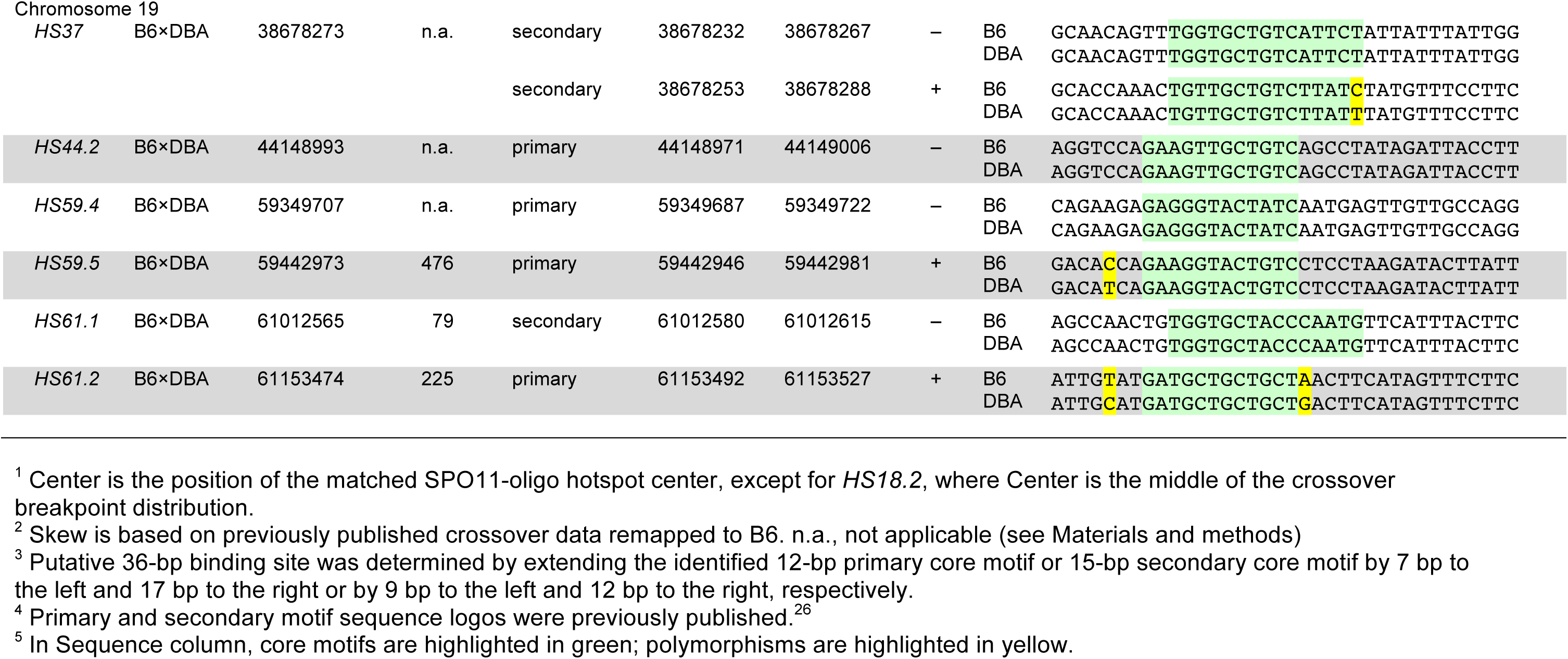
Putative PRDM9 binding sites in crossover hotspots. Using MAST^93^ to query 15 known crossover hotspots^82-88^ with previously published primary and secondary core PRDM9 motif sequences,^26^ we identified putative 36-bp PRDM9 binding sites within 250 bp of hotspot centers. Our method detected a match to the primary or secondary PRDM9 motif in 13 of the 15 crossover hotspots.

